# Improved maize reference genome with single molecule technologies

**DOI:** 10.1101/079004

**Authors:** Yinping Jiao, Paul Peluso, Jinghua Shi, Tiffany Liang, Michelle C. Stitzer, Bo Wang, Michael S. Campbell, Joshua C. Stein, Xuehong Wei, Chen-Shan Chin, Katherine Guill, Michael Regulski, Sunita Kumari, Andrew Olson, Jonathan Gent, Kevin L. Schneider, Thomas K. Wolfgruber, Michael R. May, Nathan M. Springer, Eric Antoniou, Richard McCombie, Gernot G. Presting, Michael McMullen, Jeffrey Ross-Ibarra, R. Kelly Dawe, Alex Hastie, David R. Rank, Doreen Ware

## Abstract

Complete and accurate reference genomes and annotations provide fundamental tools for characterization of genetic and functional variation. These resources facilitate elucidation of biological processes and support translation of research findings into improved and sustainable agricultural technologies. Many reference genomes for crop plants have been generated over the past decade, but these genomes are often fragmented and missing complex repeat regions. Here, we report the assembly and annotation of maize, a genetic and agricultural model species, using Single Molecule Real-Time (SMRT) sequencing and high-resolution optical mapping. Relative to the previous reference genome, our assembly features a 52-fold increase in contig length and significant improvements in the assembly of intergenic spaces and centromeres. Characterization of the repetitive portion of the genome revealed over 130,000 intact transposable elements (TEs), allowing us to identify TE lineage expansions unique to maize. Gene annotations were updated using 111,000 full-length transcripts obtained by SMRT sequencing. In addition, comparative optical mapping of two other inbreds revealed a prevalence of deletions in the low gene density region and maize lineage-specific genes.

---

Maize is the most productive and widely grown crop in the world, as well as a foundational model for genetics and genomics^1^. An accurate genome assembly for maize is critical for all forms of basic and applied research, which will enable increases in yield to feed the growing world population. The current assembly of the maize genome, based on Sanger sequencing, was first published in 2009^2^. Although this initial reference enabled rapid progress in maize genomics^3^, the original assembly is composed of more than one hundred thousand small contigs, many of which are arbitrarily ordered and oriented, significantly complicating detailed analysis of individual loci^4^ and impeding investigation of intergenic regions crucial to our understanding of phenotypic variation^5,6^ and genome evolution^7,8^.

Here we report a vastly improved *de novo* assembly and annotation of the maize reference genome (Figure 1). Based on 65X Single Molecule Real-Time sequencing we assembled the genome of the maize inbred B73 into 2,958 contigs, where half of the total assembly is made up of contigs larger than 1.2 Mb (Table 1, Extended Data Figure 1, 2a). The assembly of the long reads was then integrated with a high quality optical map (Extended Data Figure 1, Extended Data Table 1) to create a hybrid assembly consisting of 625 scaffolds (Table 1). To build chromosome-level super-scaffolds, we combined the hybrid assembly with a minimum tiling path generated from the bacterial artificial chromosomes (BACs)^9^ and a high-density genetic map^10^ (Extended Data Figure 2b). After gap-filling and error correction using short sequence reads, the total size of maize B73 RefGen_v4 pseudomolecules was 2,106 Mb. The new reference assembly has 2,522 gaps, and of which almost half (n=1,115) has optical map coverage giving an estimated mean gap length of 27 kb (Extended Data Figure 2c). The new maize B73 reference genome has 240-fold higher contiguity than the recently published short-read genome assembly of maize cultivar PH207^11^ (contig N50: 1180kb vs. 5kb).

**Figure 1.**
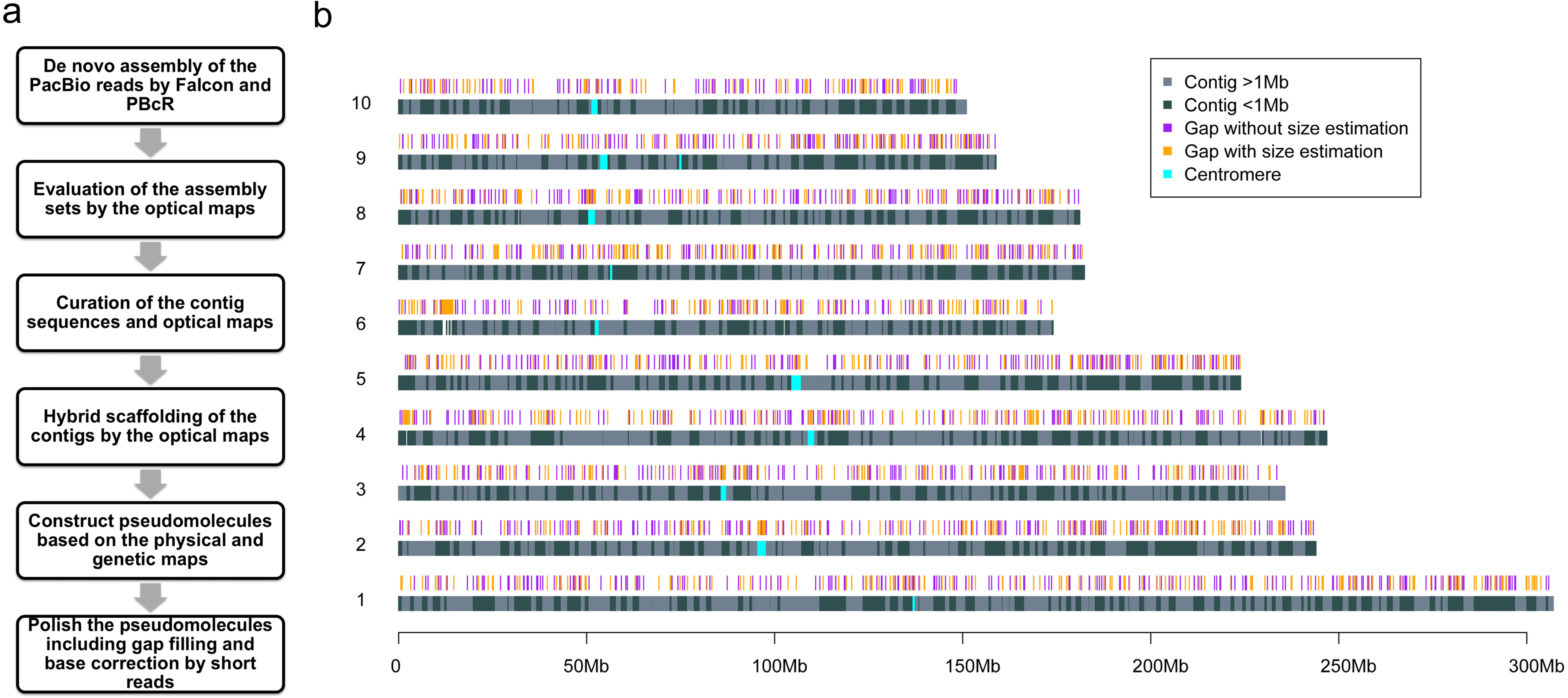
Genome assembly layout. **a)** Workflow for genome construction; **b)** Ideograms of maize B73 version 4 reference pseudomolecules. The top track shows positions of 2,522 gaps in the pseudomolecules, including 1,115 gaps whose lengths were estimated using optical genome maps (orange), while the remainder (purple) have undetermined lengths. Over half of the assembly is constituted of contigs longer than 1 Mb, which are shown as light grey bars in the bottom track.

**Table 1.** Assembly statistics of the maize B73 RefGen_v4 genome.

|  | No. of<br>contigs<br>(scaffolds) | Mean length<br>(Mb) | N50 size<br>(Mb) | Max length<br>(Mb) | Total assembly<br>length (Mb) |
| --- | --- | --- | --- | --- | --- |
| Original optical maps | 1,342 | 1.57 | 2.47 | 12.43 | 2,107 |
| Original contigs from sequence assembly | 3,303 | 0.64 | 1.04 | 5.65 | 2,105 |
| Curated optical maps | 1,356 | 1.56 | 2.47 | 12.47 | 2,114 |
| Curated contigs from sequence assembly | 2,958 | 0.71 | 1.18 | 7.26 | 2,104 |
| Optical maps in hybrid scaffolds | 1,287 | 1.62 | 2.49 | 12.47 | 2,080 |
| Contigs in hybrid scaffolds | 2,696 | 0.77 | 1.19 | 7.26 | 2,075 |
| Hybrid scaffolds | 356 | 5.97 | 9.73 | 38.53 | 2,075 |
| Hybrid scaffolds<br>+ non-scaffolded contigs | 625 | 3.45 | 9.56 | 38.53 | 2,105 |

Comparison of the new assembly to the previous BAC-based maize reference genome assembly revealed >99.9% sequence identity and a 52-fold increase in the mean contig length, with 84% of the BACs spanned by a single contig from the long reads assembly. Alignment of ChIP-seq data for the centromere-specific histone H3 (CENH3)^12^ revealed that centromeres are accurately placed and largely intact. A number of previously identified^13^ megabase-sized mis-oriented pericentromeric regions were also corrected (Extended Data Figure 3a,b). Moreover, the ends of the chromosomes are properly identified on 14 of the 20 chromosome arms based on the presence of tandem telomeric repeats and knob 180 sequences (Extended Data Figure 3a,c).

Our assembly made substantial improvements in the gene space including resolution of gaps and misassemblies and correction of order and orientation of genes. We also updated the annotation of our new assembly, resulting in consolidation of gene models (Extended Data Figure 4). Newly published full-length cDNA data^14^ improved the annotation of alternative splicing by more than doubling the number of alternative transcripts from 1.6 to 3.3 per gene (Extended Data Figure 5a) with about 70% of genes supported by the full-length transcripts. Our reference assembly also vastly improved the coverage of regulatory sequences, decreasing the number of genes exhibiting gaps in the 3 kb region(s) flanking coding sequence from 20% to <1% (Extended Data Figure 5b). The more complete sequence enabled significant improvements in the annotation of core promoter elements, especially the TATA-box, CCAAT-box, and Y patch motifs (Supplementary information). Quantitative genetic analyses have shown that polymorphism in regulatory regions explains a substantial majority of the genetic variation for many phenotypes^5,6^, suggesting that the new reference will dramatically improve our ability to identify and predict functional genetic variation.

After its divergence from *Sorghum*, the maize lineage underwent genome doubling followed by diploidization and gene loss. Previous work showed that gene loss is biased toward one of the parental genomes^2,3^, but our new assembly and annotation paint a more dramatic picture, revealing that 56% of syntenic sorghum orthologs map uniquely to the dominant maize subgenome (designated A, total size 1.16 Gb), whereas only 24% map uniquely to subgenome B (total size 0.63Gb). Gene loss in maize has primarily been considered in the context of polyploidy and functional redundancy^15^, but we found that despite its polyploidy, maize has lost a larger proportion (14%) of the 22,048 ancestral gene orthologs than any of the other four grass species evaluated to date (*Sorghum*, rice, *Brachypodium distachyon*, and *Setaria italica*, Extended Data Figure 6). Nearly one-third of these losses are specific to maize, and analysis of a restricted high-confidence set revealed enrichment for genes involved in biotic and abiotic stresses (Extended Data Table 2), e.g., NB-ARC domain disease resistance genes^16^ and the serpin protease inhibitor involved in pathogen defense and programmed cell death^17^.

Transposable elements (TEs) were first reported in maize^18^ and have since been shown to play important roles in shaping genome evolution and gene regulatory networks of many species^19^. The majority of the maize genome is derived from TEs^2,20^, and careful study of a few regions has revealed a characteristic structure of sequentially nested retrotransposons^20,21^ and the effect of deletions and recombination on retrotransposon evolution^22^. In the annotation of the original maize assembly, however, fewer than 1% of LTR retrotransposon copies were intact^23^. By applying a novel homology-independent annotation pipeline to our assembly (Extended Data Table 3), we identified 1,268Mb (130,604 copies) of structurally intact retrotransposons, of which 661 Mb (70,035 copies) are nested retrotransposon copies disrupted by the insertion of other TEs, 8.7 Mb (14,041 copies) of DNA terminal inverted repeat transposons, and 76 Mb (21,095 copies) of helitrons. To understand the evolutionary history of maize LTR retrotransposons, we also applied our annotation pipeline to the sorghum reference genome, and used reverse transcriptase protein domain sequences accessible due to the improved assembly of the internal protein coding domains of maize LTR retrotransposons to reconstruct the phylogeny of maize and sorghum LTR retrotransposon families. Despite a higher overall rate of diversification of LTR TEs in the maize lineage consistent with its larger genome size, differences in LTR retrotransposon content between genomes were primarily the result of dramatic expansion of distinct families in both lineages (Figure 2).

**Figure 2.**
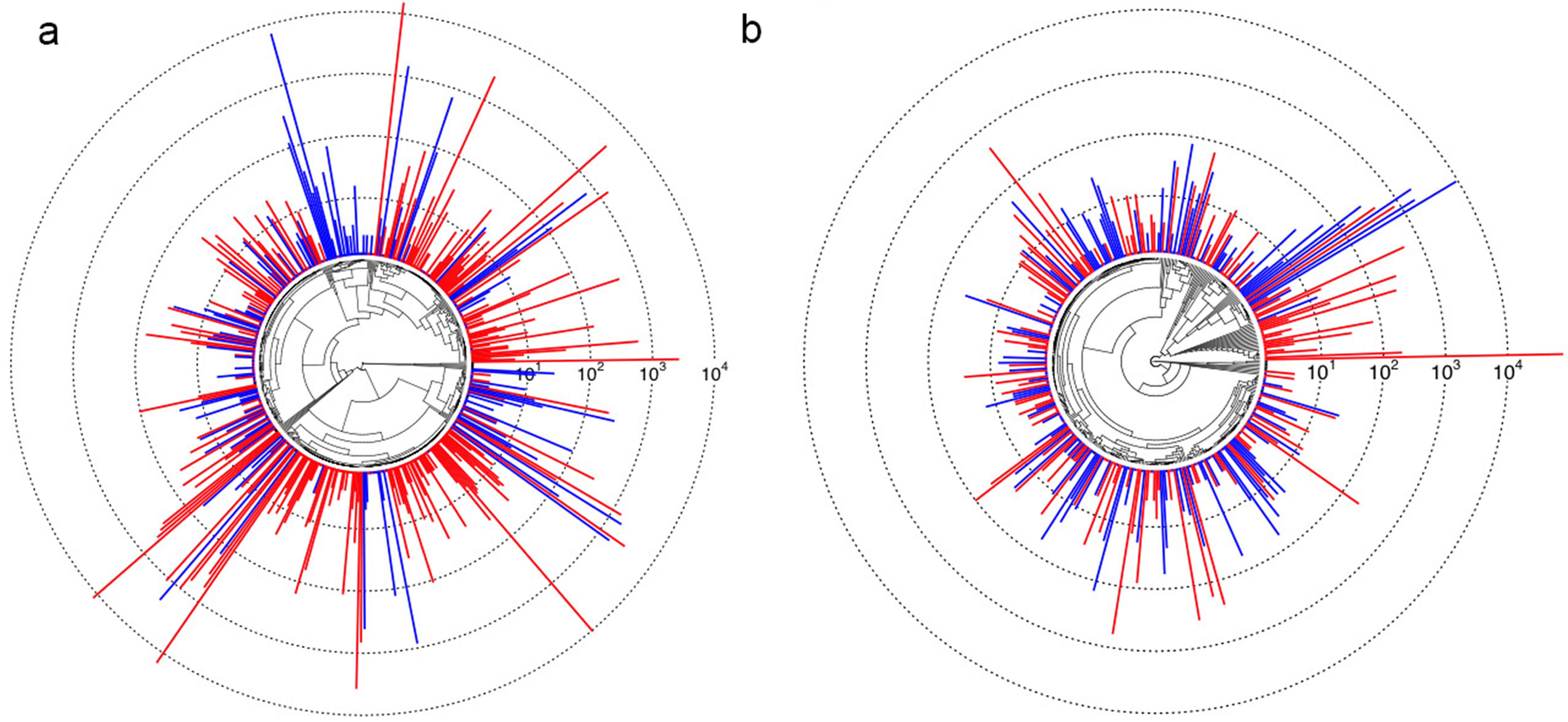
Phylogeny of maize and sorghum LTR retrotransposon families. Both **a)** Ty3/Gypsy and **b)** Ty1/Copia superfamilies are present at higher copy number in maize (red) than in sorghum (blue). Bars (log_10_-scaled) depict family copy numbers.

Maize exhibits tremendous genetic diversity^24^, and both nucleotide polymorphisms and structural variations play important roles in its phenotypic variation^8,25^. However, genome-wide patterns of structural variation in plant genomes are difficult to assess^26^, and previous efforts have relied on short-read mapping, which misses the vast majority of intergenic spaces where most rearrangements are likely to occur^8^. To investigate structural variation at a genome-wide scale, we generated optical maps (Extended Data Table 1) for two additional maize inbred lines: the tropical line Ki11, one of the founders of the maize nested association mapping (NAM) population^27^, and W22, which has served as a foundation for studies of maize genetics^28^. Due to the high degree of genomic diversity among these lines, only 32% of the assembled 2,216 Mb map of Ki11 and 39% of the 2,280 Mb W22 map could be mapped to our new B73 reference via common restriction patterns (Table 2, Figure 3A, Extended Data Figure 7). The high density of alignments across and near many of the exceedingly retrotransposon-rich centromeres reflects the comparatively low genetic diversity of most centromeres in domesticated maize^13^ and illustrates the ability of the combined optical mapping/single molecule sequencing methodology to traverse large repeat-rich regions. Within the aligned regions, approximately 32% of the Ki11 and 26% of the W22 optical maps exhibited clear evidence of structural variation, including 3,408 insertions and 3,298 deletions (Table 2). The average indel size was ∼20kb, with a range from 1kb to over 1 Mb (Figure 3B). More than 90% of the indels were unique to one inbred or the other, indicating a high level of structural diversity in maize. As short-read sequence data are available from both Ki11 and W22^8^, we analyzed 1,451 of the largest (<10kb) deletions and found that 1,083 were supported by a clear reduction in read depth (Figure 3C). The confirmed deletions occurred in regions of low gene density (4.4 genes/Mb compared to genome-wide average of 18.7 genes/Mb). One third (83/257) of the genes missing in Ki11 or W22 lack putative orthologs in all four grasses (rice, sorghum, *Brachypodium*, and *Setaria*), consistent with prior data^29^.

**Figure 3.**
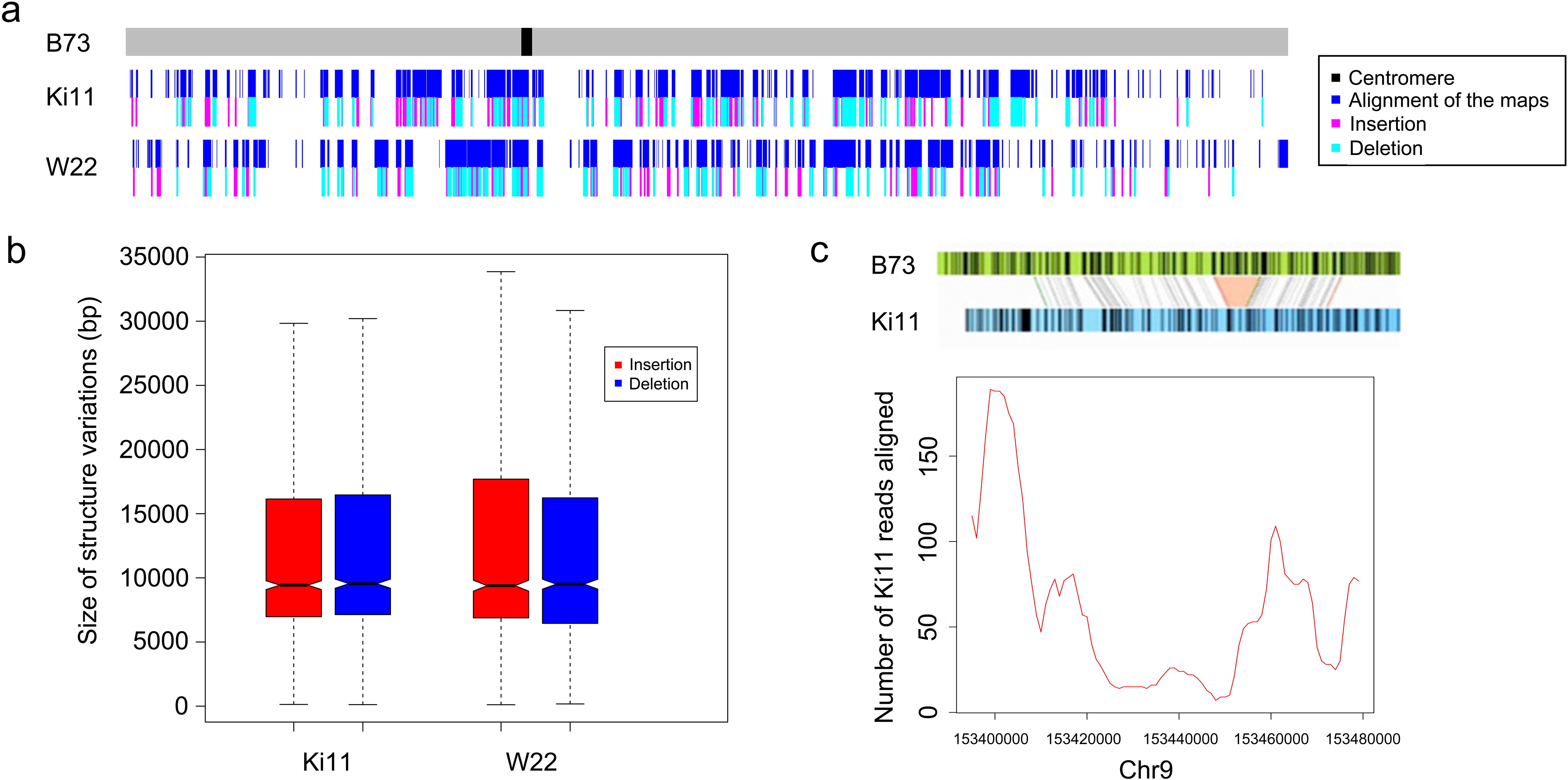
Structure variation from Ki11 and W22. **a).** Alignment and structural variation called from Ki11 and W22 optical maps on chromosome 10; **b).** Size distribution of the insertion and deletions in Ki11 and W22; **c).** Example of using short read alignment to verify a missing region mapped in Ki11.

**Table 2.**
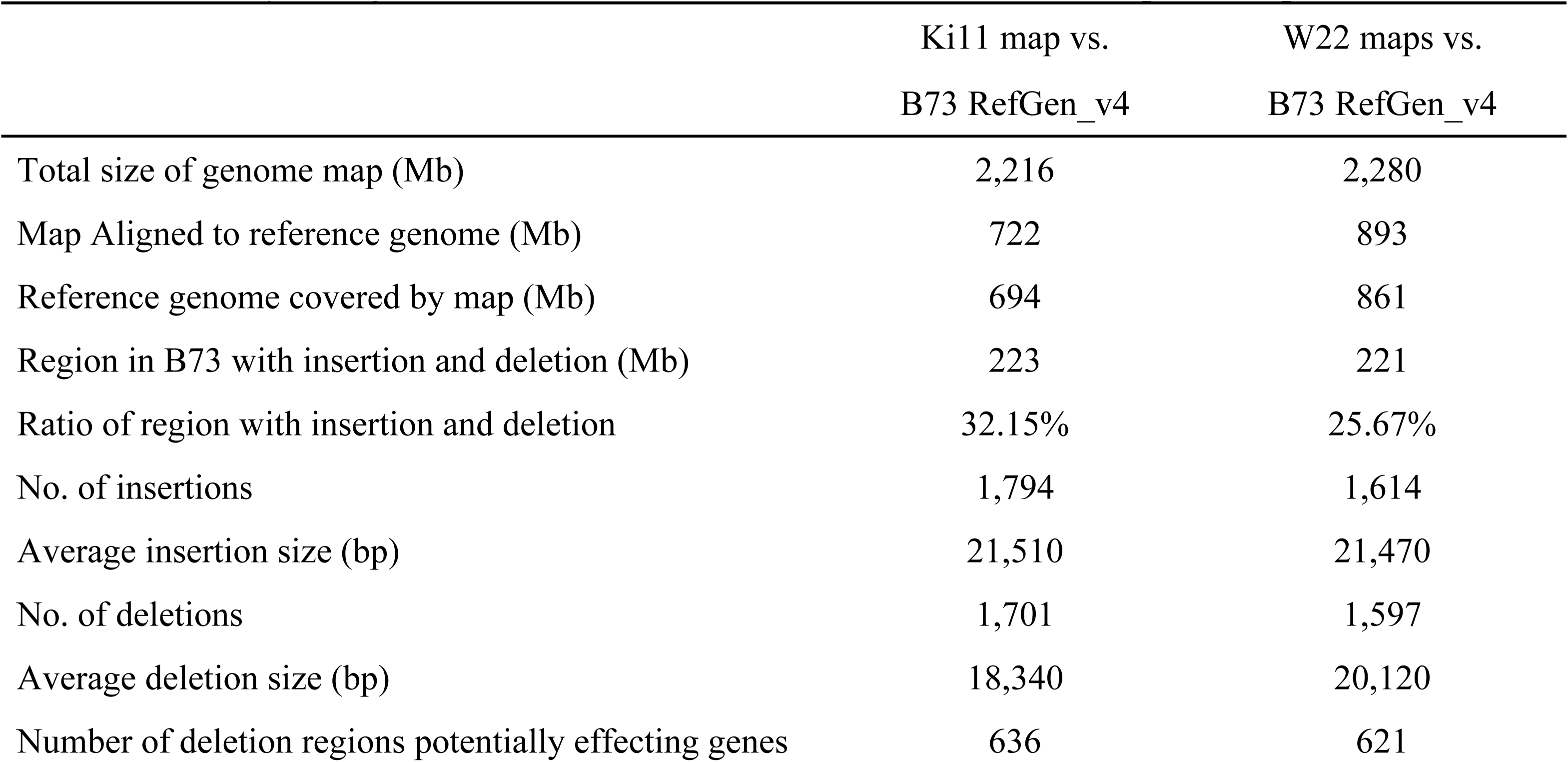
Summary of alignments and structural variations called from the optical maps of two maize lines.

Although maize is often considered to be a large-genome crop, most major food crops have even larger genomes with more complex repeat landscapes^30^. Our improved assembly of the B73 genome, generated using single molecule technologies, demonstrates that additional assemblies of other maize inbred lines and similar high-quality assemblies of other repeat-rich and large-genome plants are feasible. Additional high quality assemblies will in turn extend our understanding of the genetic diversity that forms the basis of the phenotypic diversity in maize and other economically important plants.

## Supplementary Information

## Acknowledgment

Y.J., B.W., J. S., M.C., X.W., S.K. and D.W. were supported by NSF Gramene grant IOS-1127112, NSF Cereal Gene Discovery grant 1032105, USDA-ARS CRIS 1907-21000-030-00D and NSF Plant Genome award 1238014. J.R.I. would like to acknowledge support from USDA Hatch project CA-D-PLS-2066-H and NSF Plant Genome award 1238014. R.K.D. would like to acknowledge support from NSF Plant Genome award 1444514. G.G.P. acknowledges support from NSF grant 1444624 and USDA NIFA project HAW05022-H. M.S.C would also like to acknowledge support from NSF PGRP PRFB 1523793.The authors thank Sergey Koren (National Human Genome Research Institute) for sharing genome assembly related scripts.

## Contribution

D.W. and Y.J. designed and conceived the research, M.M and K.G. prepared DNA sample for PacBio SMRT sequencing, D.R.R., P.P., E.A., and R.C. performed PacBio SMRT sequencing, B.W., J.S., R.K.D., T.L. and A.H. generated the BioNano optical genome maps, M.R. generated Illumina sequencing data, Y.J., T.L., J.S., C.C., and A.H. performed the genome assembly, J.C.S., M.S.C., X.W., B.W., Y.J., and S.K. performed gene annotation and evolutionary study, M.C.S., M.R.M., N.M.S., and J.R-I. performed transposable element analysis, J.G., J.S., R.K.D., K.L.S., T.K.W., G.G.P., and Y.J. performed the analysis of centromeres and telomeres. B.W., Y.J., J.S., T.L., A.H. and R.K.D. performed the structural variation study. X.W., J.C.S. and Y.J. contributed to the data release. Y.J., J.R-I., R.K.D., G.G.P. and D.W. wrote the paper. All authors contributed to the revision of the manuscript.

---

**Extended Data Figure 1. Summary of data generated for genome construction. a). Size distribution of single molecules for the optical maps**. A total of 150 Gb (∼60-fold coverage) of single-molecule raw data from BioNano chips was collected for map construction. The N50 of the single molecules was ∼261 kb, and the label density was 11.6 per 100 kb. After assembly, the total size of the map reached 2.12 Gb with an N50 of 2.47 Mb. **b). Length distribution of SMRT sequencing reads.** Sequencing of 212 P6-C4 SMRT cells on the PacBio platform generated ∼65-fold depth-of-coverage of the nuclear genome. Read lengths averaged 11.7 kb, with reads above 10 kb providing 53-fold depth-of-coverage. **c). The accuracy of SMRT sequencing from a representative run.** The sequencing error rate was estimated at 10% from the alignment with the maize B73 RefGen_v3 by BLASR. **d)Plot of the fraction of alignable data per run (alignable bases/total bases per chip) vs. total raw bases (per chip) for each B73 sequencing run.** As the plot shows, the trend in the data suggests that as the overall per run raw base yield increases, the fraction of alignable bases decreases. This is due to the fact that in all runs, a subset of the ZMWs will initially have more than one active sequencing enzyme in the observation field at the start of the sequencing run. A ZMW with more than one active polymerase will create un-alignable bases while the two polymerases are simultaneously synthesizing DNA and yield a ‘merged sequencing signal from two independent polymerases’. As the loading of a chips increases (yield of bases), the probability of having two or more polymerases in a single ZMW increases.

**Extended Data Figure 2. Construction of pseudomolecules. a)** Summary of the three assembly sets. **b)** How the scaffolds were ordered according to the order of the BACs. **c)** Size distribution of gaps in the pseudomolecules estimated using the optical map.

**Extended Data Figure 3. Quality assessment and comparison of the assembly in centromere and telomere regions in maize B73 RefGen_v3 and v4. a)** Quality assessment of centromere and telomere using optical genome map**. b)** Locations of centromeres on pseudomolecules defined by ChIP-seq in the B73 RefGen_v3 and v4. **c)** Telomere repeats found in the B73 RefGen_v4 pseudomolecules

**Extended Data Figure 4. Details of the gene annotation of maize B73 RefGen_V4. a)** The pipeline used to characterize high confidence gene models. **b)** Summary of B73 RefGen_v4 protein-coding gene annotation, and comparison with RefGen_v3 annotation.

**Extended Data Figure 5. Improvement of the annotation of alternative splicing and completeness of regulatory regions of maize RefGen_v4 genes. a)** Number of transcripts of each gene in v3 and v4 annotation; **b)** Percentages of genes with gaps in flanking regions in the v3 and v4 annotations.

**Extended Data Figure 6. Comparative analysis of the maize B73 RefGen_v4 genes with other grasses. a)** Species-membership in ortholog sets, giving counts and fraction (%) of ortholog sets of which each species is a member. **b)** Venn diagram showing overlap of 6,539 ortholog sets rooted in the Poaceae (true grasses) that are deficient in gene membership among five species.

**Extended Data Figure 7. Structural variation characterized from the Ki11 and W22 optical maps.**

**Extended Data Table 1. Summary of the optical maps of three maize lines.**

**Extended Data Table 2. Overrepresented protein domains in sorghum genes that lack orthologs in maize but are conserved in syntenic positions in other grasses.**

**Extended Data Table 3. Structural annotation of transposable elements.**

## Online Methods

### Data generation

#### Whole-genome sequencing using SMRT (Single Molecule Real-Time) technology

DNA samples for SMRT sequencing were prepared using maize inbred line B73 from NCRPIS (PI550473), grown at University of Missouri. Seeds of this line were deposited at NCRPIS (tracking number: PI 677128). Etiolated seedlings were grown for 4-6 days in Pro-Mix at 37°C in darkness to minimize chloroplast DNA. Batches of ∼10 g were snap-frozen in liquid nitrogen. DNA was extracted following the PacBio protocol “Preparing Arabidopsis Genomic DNA for Size-Selected ∼20 kb SMRTbell Libraries” (http://www.pacb.com/wp-content/uploads/2015/09/Shared-Protocol-Preparing-Arabidopsis-DNA-for-20-kb-SMRTbell-Libraries.pdf).

Genomic DNA was sheared to a size range of 15–40 kb using either G-tubes (Covaris^®^) or a Megarupter^®^ device (Diagenode), and enzymatically repaired and converted into SMRTbell™ template libraries as recommended by Pacific Biosciences. Briefly, hairpin adapters were ligated, after which the remaining damaged DNA fragments and those without adapters at both ends were eliminated by digestion with exonucleases. The resulting SMRTbell templates were size-selected by Blue Pippin electrophoresis (Sage Sciences) and templates ranging from 15 to 50 kb, were sequenced on a PacBio RS II instrument using P6-C4 sequencing chemistry. To acquire long reads, all data were collected as either 5- or 6-hour sequencing movies.

#### Construction of optical genome maps using the Irys system

High–molecular weight genomic DNA was isolated from three grams of young ear tissue after fixing with 2% formaldehyde. Nuclei were purified and lysed in embedded agarose as previously described^31^. DNA was labeled at Nt.BspQI sites using the IrysPrep kit. Molecules collected from BioNano chips were *de novo* assembled as previously described^32^ using “optArgument_human”.

### Genome assembly

#### *De novo* assembly of the genome sequencing data

*De novo* assembly of the long reads from SMRT Sequencing was performed using two assemblers: the Celera Assembler PBcR –MHAP pipeline^33^ and Falcon^34^ with different parameter settings. Quiver from SMRT Analysis v2.3.0 was used to polish base calling of contigs. The three independent assemblies were evaluated by aligning with the optical genome maps.

Contamination of contigs by bacterial and plasmid genomes was eliminated using the NCBI GenBank submission system^35^. Curation of the assembly, including resolution of conflicts between the contigs and the optical map and removal of redundancy at the edges of contigs, is described in the supplemental material.

#### Hybrid scaffold construction

To create hybrid scaffolds, curated sequence contigs and optical maps were aligned and merged with RefAligner^32^ (P value threshold of 1e10^−11^). These initial hybrid scaffolds were aligned again to the sequence contigs using a less stringent P value (1e10^−8^), and those contigs not previously merged were added if they aligned over 50% of their length and without overlapping previously merged contigs, thereby generating final hybrid scaffolds.

#### Pseudomolecule construction

Sequences from BACs on the physical map that were used to build the maize V3 pseudomolecules were aligned to contigs using MUMMER package^36^ with the following parameter settings: “-l(minimum length of a single match) 100 –c(the minimum length of a cluster of matches) 1000”. To only use unique hits as markers, the alignment hits were filtered with the following parameters: “-i(the minimum alignment identity) 98 –l(the minimum alignment length) 10000”. The scaffolds were then ordered and oriented into pseudochromosomes using the order of BACs as a guide. For quality control, we mapped the SNP markers from a genetic map built from an intermated maize recombinant inbred line population (Mo17×B73)^10^. Contigs with markers not located in pseudochromosomes from the physical map were placed into the AGP (A Golden Path) using the genetic map.

#### Further polishing of pseudomolecules

Raw pseudomolecules were subjected to gap filling using Pbjelly (--maxTrim=0, --minReads=2) and polished again using Quiver (SMRT Analysis v2.3.0). To increase the accuracy of the base calls, we performed two lanes of sequencing on the same genomic DNA sample (library size = 450bp) using Illumina 2500 Rapid run, which generated about 100-fold 250PE data. Reads were aligned to the assembly using BWA-mem^37^. Sequence error correction was performed with the Pilon pipeline^39^, after aligning reads with BWA-mem^37^ and parsing with SAMtools^38^, using sequence and alignment quality scores above 20.

### Annotation

For comprehensive annotation of transposable elements, we designed a structural identification pipeline incorporating several tools, including LTRharvest^40^, LTRdigest^41^, SINE-Finder^42^, MGEScan-non-LTR^43^, MITE-hunter^44^, HelitronScanner^45^, and others (details in Supplementary Information). The scripts, parameters, and intermediate files of each TE superfamily are available at https://github.com/mcstitzer/agpv4_te_annotation/tree/master/ncbi_pseudomolecule. The MAKER-P pipeline was used to annotate protein-coding genes^46^, integrating *ab initio* prediction with publicly available evidence from full-length cDNA^47^, *de novo* assembled transcripts from short read mRNA-seq^48^, Iso-Seq full-length transcripts^14^, and proteins from other species. The gene models were filtered to remove transposons and low-confidence predictions. Additional alternative transcript isoforms were obtained from the Iso-Seq data. Further details on annotations, core promoter analysis, and comparative phylogenomics are described in Supplementary Information.

### Structural Variation

Leaves were used to prepare high molecular weight DNA and optical genome maps were constructed as described above for B73. Structural variant calls were generated based on alignment to the reference map B73 V4 chromosomal assembly using the Multiple Local Alignment algorithm (RefSplit)^32^. A structural variant was identified as an alignment outlier^32,49^, defined as two well-aligned regions separated by a poorly aligned region with a large size difference between the reference genome and the map or by one or more unaligned sites, or alternately as a gap between two local alignments. A confidence score was generated by comparing the non-normalized p-values of the two well-aligned regions and the non-normalized log-likelihood ratio^50^ of the unaligned or poorly aligned region. With a confidence score threshold of 3, RefSplit is sensitive to insertions and deletions as small as 100 bp (events smaller than 1 kb are generally compound or substitution and include label changes, not just spacing differences) and other changes such as inversions and complex events which could be balanced. Insertion and deletion calls were based on an alignment outlier p-value threshold of 1× 10^−4^. Insertions or deletions that crossed gaps in the B73 pseudomolecules, or that were heterozygous in the optical genome maps, were excluded. Considering the resolution of the BioNano optical map, only insertion and deletions larger than 100 bp were used for subsequent analyses. To obtain high-confidence deletion sequences, sequencing reads from the maize HapMap2 project^8^ for Ki11 and W22 were aligned to our new B73 v4 reference genome using Bowtie2^51^. Read depth (min. mapping quality > 20) was calculated in 10-kb windows with step size of 1 kb. Windows with read depth below 10 in Ki11 and 20 in W22 (sequencing depths for Ki11 and W22 were 2.32X and 4.04X, respectively) in the deleted region were retained for further analysis.

### Data Availability

Raw reads, genome assembly sequences, and gene annotations have been deposited at NCBI under BioProject number PRJNA10769 and BioSample number SAMN04296295. PacBio whole-genome sequencing data and Illumina data were deposited in the NCBI SRA database under accessions SRX1472849 and SRX1452310, respectively. The GenBank accession number of the genome assembly and annotation is LPUQ00000000. A genome browser including genome feature tracks and ftp is available from Gramene: http://ensembl.gramene.org/Zea_mays/Info/Index

## References

1 Hake, S. & Ross-Ibarra, J. Genetic, evolutionary and plant breeding insights from the domestication of maize. eLife 4, doi:10.7554/eLife.05861 (2015).

2 Schnable, P. S. et al. The B73 maize genome: complexity, diversity, and dynamics. Science 326, 1112-1115, doi:10.1126/science.1178534 (2009).

3 Edwards, D., Batley, J. & Snowdon, R. J. Accessing complex crop genomes with next-generation sequencing. Theor Appl Genet 126, 1-11, doi:10.1007/s00122-012-1964-x (2013).

4 Fouquet, R. et al. Maize rough endosperm3 encodes an RNA splicing factor required for endosperm cell differentiation and has a nonautonomous effect on embryo development. The Plant cell 23, 4280-4297, doi:10.1105/tpc.111.092163 (2011).

5 Wallace, J. G. et al. Association mapping across numerous traits reveals patterns of functional variation in maize. PLoS Genet 10, e1004845, doi:10.1371/journal.pgen.1004845 (2014).

6 Rodgers-Melnick, E., Vera, D. L., Bass, H. W. & Buckler, E. S. Open chromatin reveals the functional maize genome. Proc Natl Acad Sci U S A 113, E3177- 3184, doi:10.1073/pnas.1525244113 (2016).

7 Hufford, M. B. et al. Comparative population genomics of maize domestication and improvement. Nat Genet 44, 808-811, doi:10.1038/ng.2309 (2012).

8 Chia, J. M. et al. Maize HapMap2 identifies extant variation from a genome in flux. Nat Genet 44, 803-807, doi:10.1038/ng.2313 (2012).

9 Wei, F. et al. The physical and genetic framework of the maize B73 genome. PLoS Genet 5, e1000715, doi:10.1371/journal.pgen.1000715 (2009).

10 Ganal, M. W. et al. A large maize (Zea mays L.) SNP genotyping array: development and germplasm genotyping, and genetic mapping to compare with the B73 reference genome. PLoS One 6, e28334, doi:10.1371/journal.pone.0028334 (2011).

11 Hirsch, C. N. et al. Draft Assembly of Elite Inbred Line PH207 Provides Insights into Genomic and Transcriptome Diversity in Maize. The Plant cell 28, 2700-2714, doi:10.1105/tpc.16.00353 (2016).

12 Gent, J. I., Wang, K., Jiang, J. & Dawe, R. K. Stable Patterns of CENH3 Occupancy Through Maize Lineages Containing Genetically Similar Centromeres. Genetics 200, 1105-1116, doi:10.1534/genetics.115.177360 (2015).

13 Schneider, K. L., Xie, Z., Wolfgruber, T. K. & Presting, G. G. Inbreeding drives maize centromere evolution. Proc Natl Acad Sci U S A 113, E987-996, doi:10.1073/pnas.1522008113 (2016).

14 Wang, B. et al. Unveiling the complexity of the maize transcriptome by single-molecule long-read sequencing. Nature communications 7, 11708, doi:10.1038/ncomms11708 (2016).

15 Schnable, J. C., Springer, N. M. & Freeling, M. Differentiation of the maize subgenomes by genome dominance and both ancient and ongoing gene loss. Proc Natl Acad Sci U S A 108, 4069-4074, doi:10.1073/pnas.1101368108 (2011).

16 McHale, L., Tan, X., Koehl, P. & Michelmore, R. W. Plant NBS-LRR proteins: adaptable guards. Genome biology 7, 212, doi:10.1186/gb-2006-7-4-212 (2006).

17 Fluhr, R., Lampl, N. & Roberts, T. H. Serpin protease inhibitors in plant biology. Physiologia plantarum 145, 95-102, doi:10.1111/j.1399-3054.2011.01540.x (2012).

18 McClintock, B. The origin and behavior of mutable loci in maize. Proc Natl Acad Sci U S A 36, 344–355 (1950).

19 Slotkin, R. K. & Martienssen, R. Transposable elements and the epigenetic regulation of the genome. Nature reviews. Genetics 8, 272-285, doi:10.1038/nrg2072 (2007).

20 SanMiguel, P. et al. Nested retrotransposons in the intergenic regions of the maize genome. Science 274, 765–768 (1996).

21 Brunner, S., Fengler, K., Morgante, M., Tingey, S. & Rafalski, A. Evolution of DNA sequence nonhomologies among maize inbreds. The Plant cell 17, 343– 360 (2005).

22 Sharma, A., Schneider, K. L. & Presting, G. G. Sustained retrotransposition is mediated by nucleotide deletions and interelement recombinations. Proceedings of the National Academy of Sciences 105, 15470–15474 (2008).

23 Baucom, R. S. et al. Exceptional diversity, non-random distribution, and rapid evolution of retroelements in the B73 maize genome. PLoS Genet 5, e1000732 (2009).

24 Buckler, E. S., Gaut, B. S. & McMullen, M. D. Molecular and functional diversity of maize. Current opinion in plant biology 9, 172-176, doi:10.1016/j.pbi.2006.01.013 (2006).

25 Dooner, H. K. & He, L. Maize genome structure variation: interplay between retrotransposon polymorphisms and genic recombination. The Plant cell 20, 249-258, doi:10.1105/tpc.107.057596 (2008).

26 Saxena, R. K., Edwards, D. & Varshney, R. K. Structural variations in plant genomes. Briefings in functional genomics 13, 296-307, doi:10.1093/bfgp/elu016 (2014).

27 McMullen, M. D. et al. Genetic properties of the maize nested association mapping population. Science 325, 737-740, doi:10.1126/science.1174320 (2009).

28 Strable, J. & Scanlon, M. J. Maize (Zea mays): a model organism for basic and applied research in plant biology. Cold Spring Harbor protocols 2009, pdb emo132, doi:10.1101/pdb.emo132 (2009).

29 Swanson-Wagner, R. A. et al. Pervasive gene content variation and copy number variation in maize and its undomesticated progenitor. Genome research 20, 1689-1699, doi:10.1101/gr.109165.110 (2010).

30 Morrell, P. L., Buckler, E. S. & Ross-Ibarra, J. Crop genomics: advances and applications. Nature reviews. Genetics 13, 85-96, doi:10.1038/nrg3097 (2011).

## References

31 VanBuren, R. et al. Single-molecule sequencing of the desiccation-tolerant grass Oropetium thomaeum. Nature 527, 508-511, doi:10.1038/nature15714 (2015).

32 Pendleton, M. et al. Assembly and diploid architecture of an individual human genome via single-molecule technologies. Nat Methods 12, 780-786, doi:10.1038/nmeth.3454 (2015).

33 Berlin, K. et al. Assembling large genomes with single-molecule sequencing and locality-sensitive hashing. Nature biotechnology 33, 623-630, doi:10.1038/nbt.3238 (2015).

34 Chin, C. S. et al. Phased diploid genome assembly with single-molecule real-time sequencing. Nat Methods 13, 1050-1054, doi:10.1038/nmeth.4035 (2016).

35 Clark, K., Karsch-Mizrachi, I., Lipman, D. J., Ostell, J. & Sayers, E. W. GenBank. Nucleic Acids Res 44, D67-72, doi:10.1093/nar/gkv1276 (2016).

36 Kurtz, S. et al. Versatile and open software for comparing large genomes. Genome biology 5, R12, doi:10.1186/gb-2004-5-2-r12 (2004).

37 Li, H. & Durbin, R. Fast and accurate short read alignment with Burrows-Wheeler transform. Bioinformatics 25, 1754-1760, doi:10.1093/bioinformatics/btp324 (2009).

38 Li, H. et al. The Sequence Alignment/Map format and SAMtools. Bioinformatics 25, 2078-2079, doi:10.1093/bioinformatics/btp352 (2009).

39 Walker, B. J. et al. Pilon: an integrated tool for comprehensive microbial variant detection and genome assembly improvement. PLoS One 9, e112963, doi:10.1371/journal.pone.0112963 (2014).

40 Ellinghaus, D., Kurtz, S. & Willhoeft, U. LTRharvest, an efficient and flexible software for de novo detection of LTR retrotransposons. BMC bioinformatics 9, 18, doi:10.1186/1471-2105-9-18 (2008).

41 Steinbiss, S., Willhoeft, U., Gremme, G. & Kurtz, S. Fine-grained annotation and classification of de novo predicted LTR retrotransposons. Nucleic Acids Res 37, 7002-7013, doi:10.1093/nar/gkp759 (2009).

42 Wenke, T. et al. Targeted identification of short interspersed nuclear element families shows their widespread existence and extreme heterogeneity in plant genomes. The Plant cell 23, 3117-3128, doi:10.1105/tpc.111.088682 (2011).

43 Rho, M. & Tang, H. MGEScan-non-LTR: computational identification and classification of autonomous non-LTR retrotransposons in eukaryotic genomes. Nucleic Acids Res 37, e143, doi:10.1093/nar/gkp752 (2009).

44 Han, Y. & Wessler, S. R. MITE-Hunter: a program for discovering miniature inverted-repeat transposable elements from genomic sequences. Nucleic Acids Res 38, e199, doi:10.1093/nar/gkq862 (2010).

45 Xiong, W., He, L., Lai, J., Dooner, H. K. & Du, C. HelitronScanner uncovers a large overlooked cache of Helitron transposons in many plant genomes. Proc Natl Acad Sci U S A 111, 10263-10268, doi:10.1073/pnas.1410068111 (2014).

46 Campbell, M. S. et al. MAKER-P: a tool kit for the rapid creation, management, and quality control of plant genome annotations. Plant Physiol 164, 513-524, doi:10.1104/pp.113.230144 (2014).

47 Soderlund, C. et al. Sequencing, mapping, and analysis of 27,455 maize full-length cDNAs. PLoS Genet 5, e1000740, doi:10.1371/journal.pgen.1000740 (2009).

48 Law, M. et al. Automated update, revision, and quality control of the maize genome annotations using MAKER-P improves the B73 RefGen_v3 gene models and identifies new genes. Plant Physiol 167, 25-39, doi:10.1104/pp.114.245027 (2015).

49 Mostovoy, Y. et al. A hybrid approach for de novo human genome sequence assembly and phasing. Nat Methods, doi:10.1038/nmeth.3865 (2016).

50 Cao, H. et al. Rapid detection of structural variation in a human genome using nanochannel-based genome mapping technology. GigaScience 3, doi:10.1186/2047-217X-3-34 (2014).

51 Langmead, B. & Salzberg, S. L. Fast gapped-read alignment with Bowtie 2. Nat Methods 9, 357-359, doi:10.1038/nmeth.1923 (2012).

